# CAMK2D causes heart failure in RBM20 cardiomyopathy

**DOI:** 10.1101/2025.05.23.655557

**Authors:** Maarten MG van den Hoogenhof, Javier Duran, Thiago Britto-Borges, Elena Kemmling, Laura Konrad, David C Lennermann, Joshua Hartmann, Laura Schraft, Julia Kornienko, Theresa Bock, Marcus Krüger, Christoph Dieterich, Lars M Steinmetz, Matthias Dewenter, Johannes Backs

## Abstract

Although heart disease can arise from different etiologies, current treatment is not tailored to the different underlying causes but is rather a one-size-fits-all approach. Importantly, not all patients benefit from this treatment regimen, which means the number needed to treat is very high.

Moreover, this makes clinical trials large and costly, limiting clinical translation. Thus, there is a high medical need to develop a first etiology-specific therapy. Mutations in RBM20, a splicing factor that targets multiple pivotal cardiac genes including *TTN* and *CAMK2D,* cause a clinically aggressive form of dilated cardiomyopathy (DCM) with a high risk of malignant ventricular arrhythmias. Here, we hypothesized that *CAMK2D* is the heart disease-causing target of RBM20. We crossed *Camk2d*- to *Rbm20*-deficient mice and found that double knockout (DKO) mice were protected from heart failure and sudden cardiac death. Phosphorylation of multiple CAMK2D targets was increased in *Rbm20*-deficient mouse hearts, which was reverted in DKO hearts, confirming that CAMK2D is not only misspliced but also overactivated. AAV9-mediated re-expression of single CAMK2D splice variants in DKO mice reintroduced cardiac dysfunction irrespective of the splice variant, unmasking that overactivation rather than missplicing underlies the detrimental phenotype. To test whether heart failure could pharmacologically be reversed, we treated heterozygous *Rbm20*-R636Q knock-in (KI) mice with hesperadin, a potent CAMK2 inhibitor, which rescued both cardiac function and ventricular geometry. These data demonstrate that overactivation of CAMK2D underlies heart failure in RBM20 cardiomyopathy. Pharmacological inhibition of CAMK2D could therefore become the first cause-directed DCM therapy.

## INTRODUCTION

Heart disease is the most frequent cause of death and is often caused by dilated cardiomyopathy (DCM) [1]. DCM is defined by left ventricular or biventricular systolic dysfunction and dilation that are not explained by abnormal loading conditions or coronary artery disease [2]. Prevalence of DCM is approximately 1:250, of which ∼30-50% of cases are familial [3–5]. Mutations that are causal for familial DCM can be found in a variety of genes, including *RBM20*, which accounts for approximately 3% of familial DCM cases [3]. Sadly, despite the specific etiologies, there is currently no specific DCM therapy. In stark contrast, the development of tailored therapies in oncology has revolutionized the clinical care of cancer patients [6]. In the cardiac field, the development of mavacamten for hypertrophic cardiomyopathy (HCM) has demonstrated that the treatment of the specific underlying pathophysiology can also improve the outcome of cardiology patients [7]. RBM20, or RNA binding motif protein 20, is a cardiac enriched splice factor. RBM20 mutations lead to a severe form of DCM characterized by an early onset of disease and a high burden of malignant arrhythmias causing sudden cardiac death, and is generally termed RBM20 cardiomyopathy [8]. RBM20 is involved in proper splicing of a multitude of cardiac genes, including *TTN*, *RYR2*, *CAMK2D*, and *LDB3* [9, 10]. Loss of functional RBM20 generally leads to inclusion of exons, and RBM20 is therefore regarded as a splicing repressor [11]. Many of the studies on RBM20 have focused on TTN, which is misspliced into a giant and heavily compliant isoform, due to the inclusion of exons between exon 50 and 219, termed N2BA-Giant [10, 11]. It has been hypothesized and shown that an increase in this isoform through downregulation of RBM20 might be beneficial, especially in the setting of diastolic dysfunction [12, 13]. However, an increase in this TTN isoform could potentially impact and decrease systolic function. Another direct target of RBM20 is the multifunctional calcium/calmodulin dependent kinase II delta (*CAMK2D*). In the heart, *CAMK2D* is known to have at least 11 different splice isoforms, of which CAMK2D-9 is the most expressed isoform in the human heart [14]. The different splice isoforms of CAMK2D have overlapping, but also distinct functions, that are currently not completely unraveled [15]. Increased activation of CAMK2D contributes to a variety of pathological cardiac conditions, including ischemic heart disease and pressure-overload induced heart failure, as it is seen in chronic arterial hypertension. However, it is unknown whether a specific splice variant or several variants are mainly responsible for these detrimental effects. CAMK2D works in heteromultimeres, and can include different splice isoforms in different ratios. It is hypothesized that the ratio of these splice isoforms determines the preference of the enzyme for localization in different cellular compartments or for the phosphorylation of different proteins [16]. For example, CAMK2D-B is mostly nuclear and is known for its involvement in transcription regulation [17, 18], while CAMK2D-C is mostly cytoplasmic and is known for its involvement in calcium handling and mitochondrial regulation [19, 20]. In RBM20 cardiomyopathy, CAMK2D is misspliced and activated, but its contribution to the molecular mechanisms that underlie RBM20 cardiomyopathy is unknown. Furthermore, recent findings suggest that the clinical phenotype of RBM20 cardiomyopathy is not solely caused by a loss of splicing, but that mutations in the RS-region of RBM20 lead to cytoplasmic ribonucleoprotein granule formation, which is an additional detrimental mechanism [21, 22]. Specifically, mutations in the RS domain (especially in the RSRSP stretch) of RBM20 disrupt its interaction with the nuclear import receptor TNPO3, resulting in mislocalization and cytoplasmic retention [23]. This mechanism is supported by the observation that pig, mouse, and hiPSC with mutations in the RS region of RBM20 have a more severe phenotype than *Rbm20* KO models, highlighting the pathological significance of RBM20 mislocalization alongside its splicing dysfunction [21, 22, 24, 25]. Here, we hypothesised that CAMK2D is a critical target of RBM20, and generated *Rbm20*/*Camk2d* double knockout (DKO) mice. We characterized these mice functionally and molecularly, and showed that DKO mice have similar splicing abnormalities in RBM20 targets, but were protected against cardiac dysfunction and lethal arrhythmias. We further show that this effect is dependent on CAMK2D overactivation, but independent on splice isoform expression. Lastly, we show in a recently established mouse model of RBM20 cardiomyopathy, carrying a patient mutation in the RS domain and characterized by cytoplasmic ribonucleoprotein granule formation, that pharmacological CAMK2D inhibition reverses heart failure. These data open the door to a first specific and cause-directed therapy for DCM patients due to RBM20 mutations.

## RESULTS

### CAMK2D is misspliced in RBM20 cardiomyopathy

We hypothesized that CAMK2D is a critical target of RBM20, and therefore we characterized CAMK2D splicing in more detail in multiple models of RBM20 cardiomyopathy. We made use of existing RNA-sequencing datasets from mouse models of RBM20 cardiomyopathy [26, 27], as well as hiPSC-CM with disease-causing mutations in the *RBM20*-gene [28]. We first used *Rbm20* KO mice, and identified splice junction reads that were specific to the different cardiac CAMK2D splice variants CAMK2D-A, CAMK2D-B, CAMK2D-C, CAMK2D-4, and CAMK2D-9, and corrected them for total expression of CAMK2D (exon composition of the different splice variants is depicted in Figure 1A). We found that in wildtype (WT) mouse hearts CAMK2D-9 was the most prevalent isoform, followed by CAMK2D-C and CAMK2D-B which is in line with recent insight, and loss of RBM20 led to an almost complete switch to CAMK2D-A and an increase in CAMK2D-4 (Figure 1B) [14]. Two mouse models with a mutation in *Rbm20* (P635L and R636Q) showed similar missplicing of CAMK2D (Figure 1C). We then investigated CAMK2D splicing in publicly available RNA-sequencing data from a human model of RBM20 cardiomyopathy, namely hiPSC with the P633L and R634Q mutation in *RBM20* [28]. Interestingly, we similarly observed missplicing of CAMK2D, but the isoforms expressed were different. In mice, CAMK2D-A and CAMK2D-4 were increased, but in humans we observed an increase in CAMK2D-9 (Figure 1D). This difference is due to the lack of expression of exon 15 in humans, which is included in CAMK2D-A and CAMK2D-4, but not in CAMK2D-9 [10, 29]. Overall, this data shows that loss of functional RBM20 leads to missplicing of *CAMK2D*, albeit with differences between species.

**Figure 1.**
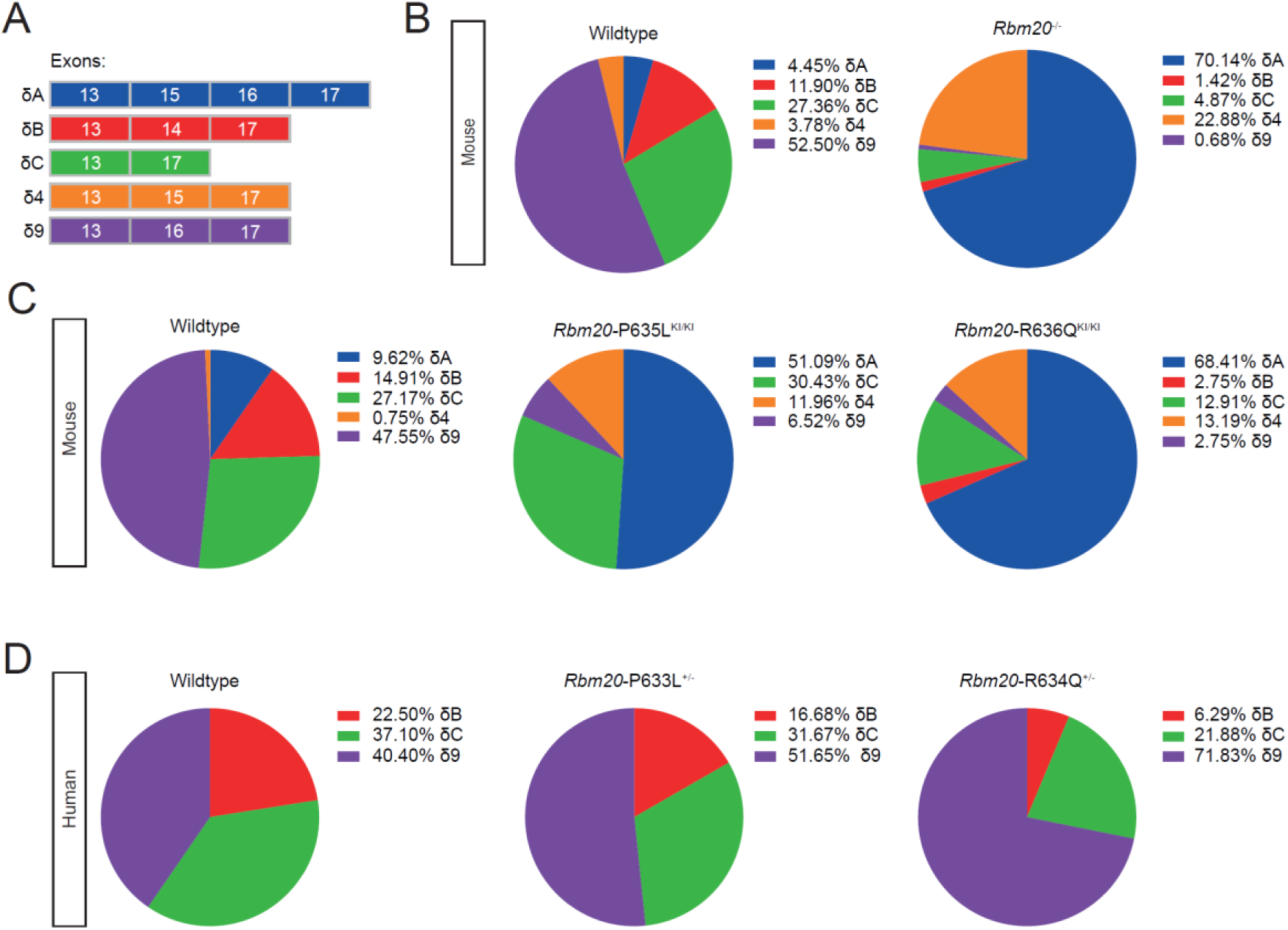
*Camk2d* is misspliced after loss of functional RBM20. A. Overview of exons between exon 13 and 17 that are included in the different *Camk2d* splice variants. B. Pie charts of the cardiac *Camk2d* splice variants in WT and *Rbm20* KO mouse hearts. C. Pie charts of the cardiac *Camk2d* splice variants in WT, *Rbm20*-P635L, and *Rbm20*-R636Q KI mouse hearts. D. Pie charts of the cardiac *CAMK2D* splice variants in WT, *RBM20*-P633L and *RBM20*-R634Q mutant hiPSC-CM.

### *Rbm20*/*Camk2d* DKO mice are protected against cardiac dysfunction

To investigate the contribution of the RBM20 splicing target CAMK2D to RBM20 cardiomyopathy, we generated homozygous *Rbm20*/*Camk2d* DKO mice (Figure 2). Histochemistry showed enlarged atria upon loss of *Rbm20* in *Rbm20* KO and DKO, but no other observable morphological changes (Figure 2A). *Rbm20* expression was abolished in both the *Rbm20* KO and the DKO mice (Figure 2B). CAMK2D protein was absent from the hearts of *Camk2d* KO and DKO mice (Figure 2C). Next, we performed cardiac echocardiography and found that *Rbm20* KO mice presented with decreased cardiac function, while DKO mice were protected against cardiac dysfunction (Figure 2D, Supp Figure 1).

**Figure 2.**
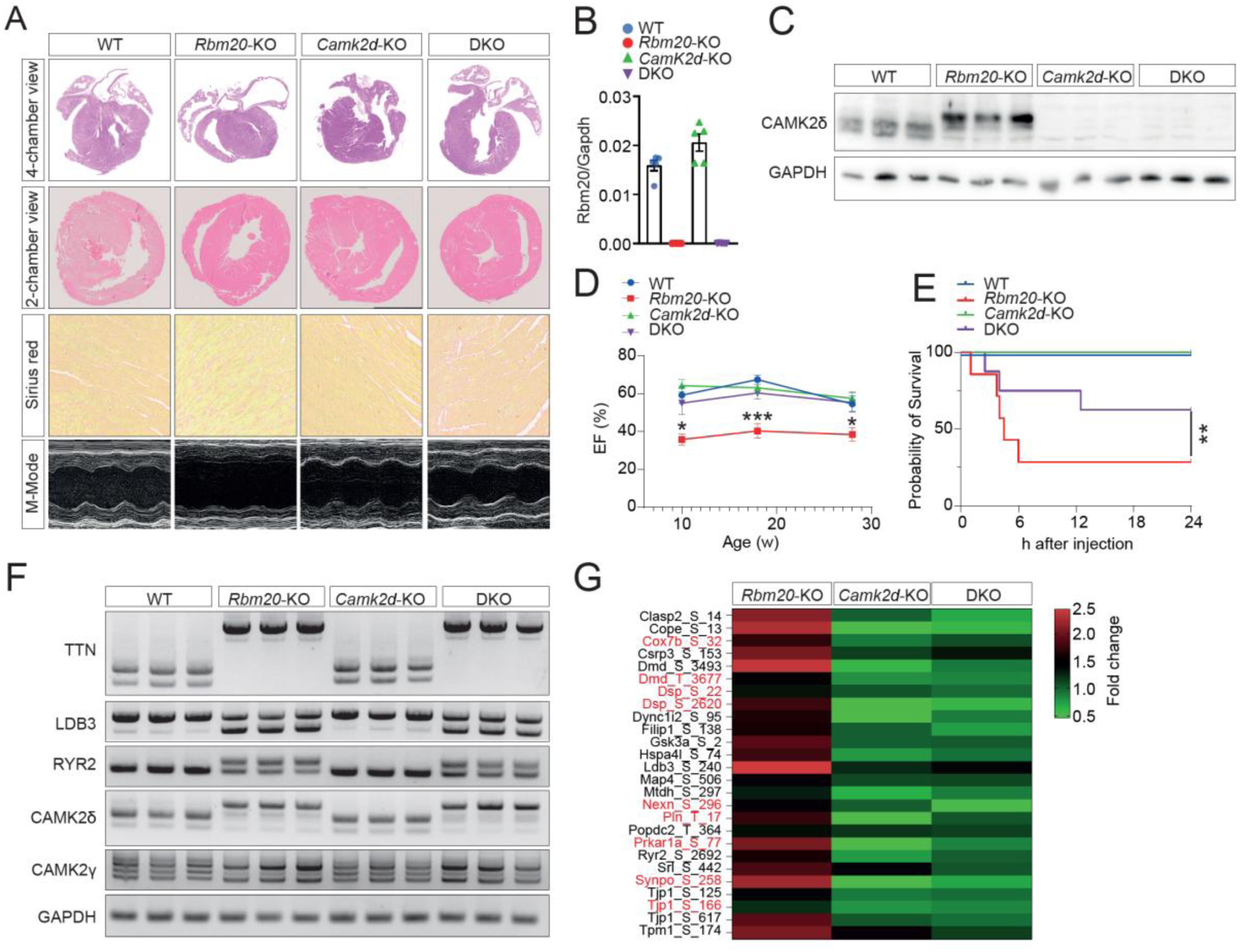
*Rbm20*/*Camk2d* DKO mice are protected from cardiac dysfunction and sudden cardiac death. A. H&E and Picrosirius Red staining of hearts of WT, *Rbm20* KO, *Camk2d* KO, and *Rbm20/Camk2d* DKO mice. B. qPCR of *Rbm20*. C. Western blotting of CAMK2D in left ventricular tissue. D. Cardiac function measured by echocardiography. Significance was tested using one-way ANOVA. E. Survival after arrhythmia induction. Significance was tested using a Log-rank (Mantel-Cox) test. F. RT-PCR of RBM20 splicing targets. G. Heatmap of differentially phosphorylated proteins that are increased in the *Rbm20* KO, and normalized in the DKO. Proteins in red are known/predicted CAMK2 phosphorylation targets.

Diastolic function was similarly rescued in DKO mice (Supp Figure 2). In addition, DKO mice were partly protected against sudden cardiac death after arrhythmia induction (Figure 2E). We then investigated known RBM20 splicing targets, and found that *Ttn*, *Ldb3*, *Ryr2*, and *Camk2g* were similarly misspliced in DKO as in *Rbm20* KO mouse hearts (Figure 2F). With regards to *Camk2d*, we observed similar missplicing on the transcript level, although protein was completely absent from the DKO hearts, which is due to a second transcriptional start site in exon 8 which does not result in a protein (Figure 2C-F) [30]. Interestingly, apart from missplicing of *Camk2d*, we also observed an increase in total CAMK2D protein, which correlates with increased CAMK2D activity (Figure 2C).

Phosphoproteomics on the hearts of WT, *Rbm20* KO, *Camk2d* KO, and DKO mice revealed increased phosphorylation of 136 protein residues in *Rbm20* KO hearts, of which 27 were normalized in DKO hearts (Figure 2G, Supp Figure 3). Of these 27 residues, 9 were known or predicted targets of CAMK2, including PLN-T17 and PRKAR1A-S77. This suggests that CAMK2 activity is increased in the hearts of *Rbm20* KO hearts, and this is alleviated in the hearts of DKO mice. Bulk RNA-sequencing on the hearts of WT, *Rbm20* KO, *Camk2d* KO, and DKO mice revealed a subset of genes (cluster 2) that reverted back to normal levels (Supp Figure 4). Overall, these data show that *Rbm20* KO mice present with cardiac dysfunction and increased susceptibility to sudden cardiac death, which is alleviated by genetic deletion of *Camk2d*.

### Re-expression of single *Camk2d* splice variants in DKO mice reintroduces cardiac dysfunction

To investigate which cardiac splice variant(s) of *Camk2d* are required for the cardiomyopathic phenotype in RBM20 cardiomyopathy, we used AAV9-mediated overexpression of the cardiac *Camk2d* splice variants CAMK2D-A, CAMK2D-B, CAMK2D-C, CAMK2D-4, and CAMK2D-9 (depicted as δA, δB, δC, δ4, and δ9) in the hearts of DKO mice. We aimed to re-express the different splice variants to similar levels as seen in the *Rbm20* KO. We injected mice with 2 x 10^12^ viral genomes of AAV9 encoding N-terminal flag-tagged splice variants of *Camk2d*, performed echocardiography and harvested the organs 4 weeks later (Figure 3A). Immunoblotting revealed specific expression of flag-tagged CAMK2D in the AAV9-injected mice, and that CAMK2D expression was restored to physiological levels in the AAV9-injected mice (Figure 3B). Cardiac function was impaired in *Rbm20* KO mice, and DKO mice injected with the control AAV9-Luciferase were, as expected, protected against cardiac dysfunction (Figure 3C). Interestingly, cardiac function was markedly reduced in all DKO mice with re-expression of a *Camk2d* splice variant, regardless of which splice variant was reintroduced (Figure 3C). Since this suggests that there is at least a partial overlap in function between the different splice isoforms, we investigated cellular localization of the different CAMK2D splice isoforms *in vivo* by doing cellular fractionation of WT and *Rbm20* KO hearts. Surprisingly, even though *Camk2d* isoform expression is markedly different between WT and *Rbm20* KO mice, we found almost all CAMK2D in the nuclear fraction of both groups, regardless of which isoform was expressed (Supp Figure 5). This suggests that the shift in CAMK2D splice isoform expression has only limited effect on localization, and potentially function, of the enzyme, at least when ectopically expressed *in vivo*. Expression of the cardiac stress markers *Anf* and *Bnp* was increased in the hearts of all mice, except WT, and we did not observe any morphological differences between the groups (Figure 3D, Supp Figure 6). Overall, these results show that re-expression of CAMK2D in DKO mouse hearts is sufficient to re-introduce cardiac dysfunction, regardless of which splice variant is re- introduced. Moreover, this suggests that (over)activation of CAMK2D, and not missplicing, underlies the disease phenotype in *Rbm20* KO mice.

**Figure 3.**
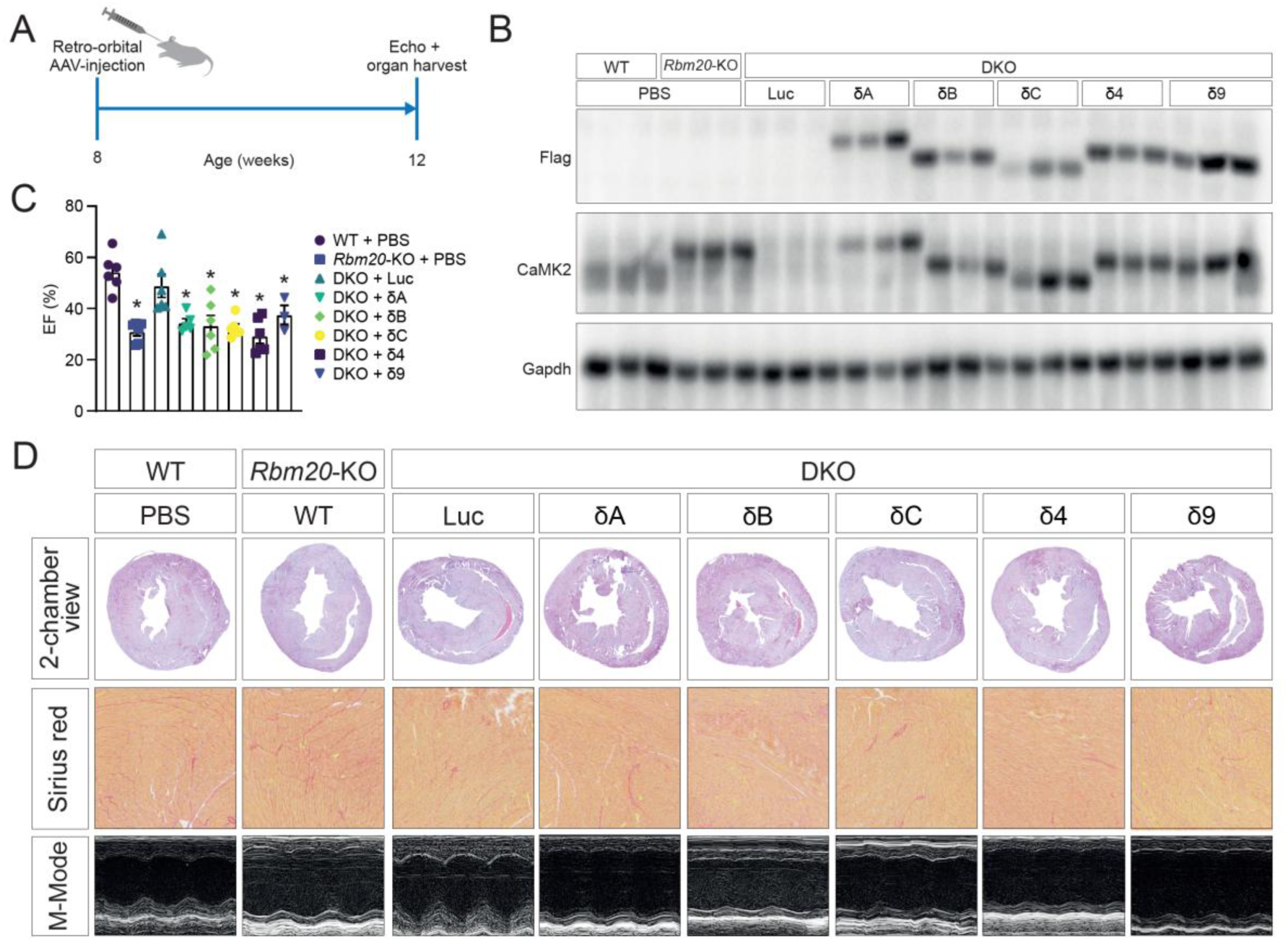
Re-expression of CAMK2D splice variants reintroduces cardiac dysfunction in *Rbm20/Camk2d* DKO mice. A. Experimental set-up. B. Western blotting of CAMK2D and Flag in left ventricular tissue. C. Cardiac function as measured by echocardiography. Significance was tested with a one-way ANOVA. D. H&E and Picrosirius Red staining.

### Hesperadin treatment ameliorates cardiac dysfunction in *Rbm20*-R636Q KI mice

To evaluate the therapeutic potential of CAMK2D inhibition in RBM20 cardiomyopathy, we used hesperadin, a novel CAMK2 inhibitor, in *Rbm20*-R636Q KI mice, which harbor a human disease-causing mutation that leads to cytoplasmic granule formation. We treated heterozygous *Rbm20*- R636Q KI mice for 4 weeks with either vehicle (DMSO) or hesperadin (2.5 mg/kg or 5 mg/kg) (Figure 4A) [31]. At 8 weeks old, heterozygous *Rbm20*-R636Q KI mice presented with decreased cardiac function and increased diastolic volume as compared to WT mice (Figure 4B). Hesperadin treatment completely restored cardiac function, both at 2 weeks and at 4 weeks after begin of treatment. In addition, treatment reduced diastolic volume in the *Rbm20*-R636Q KI mouse hearts suggesting beneficial effects on structural remodelling (Figure 4B). Histochemistry revealed no gross morphological changes between vehicle and hesperadin treated mice (Figure 4C). *Rbm20* mRNA expression was not affected by hesperadin treatment (Figure 4D). Furthermore, we used immunohistochemistry to investigate whether hesperadin treatment affected cytoplasmic granule formation, but granule formation was unaffected, pointing to a process downstream of cytoplasmic granule formation that leads to CAMK2D overactivation (Figure 4E).

**Figure 4.**
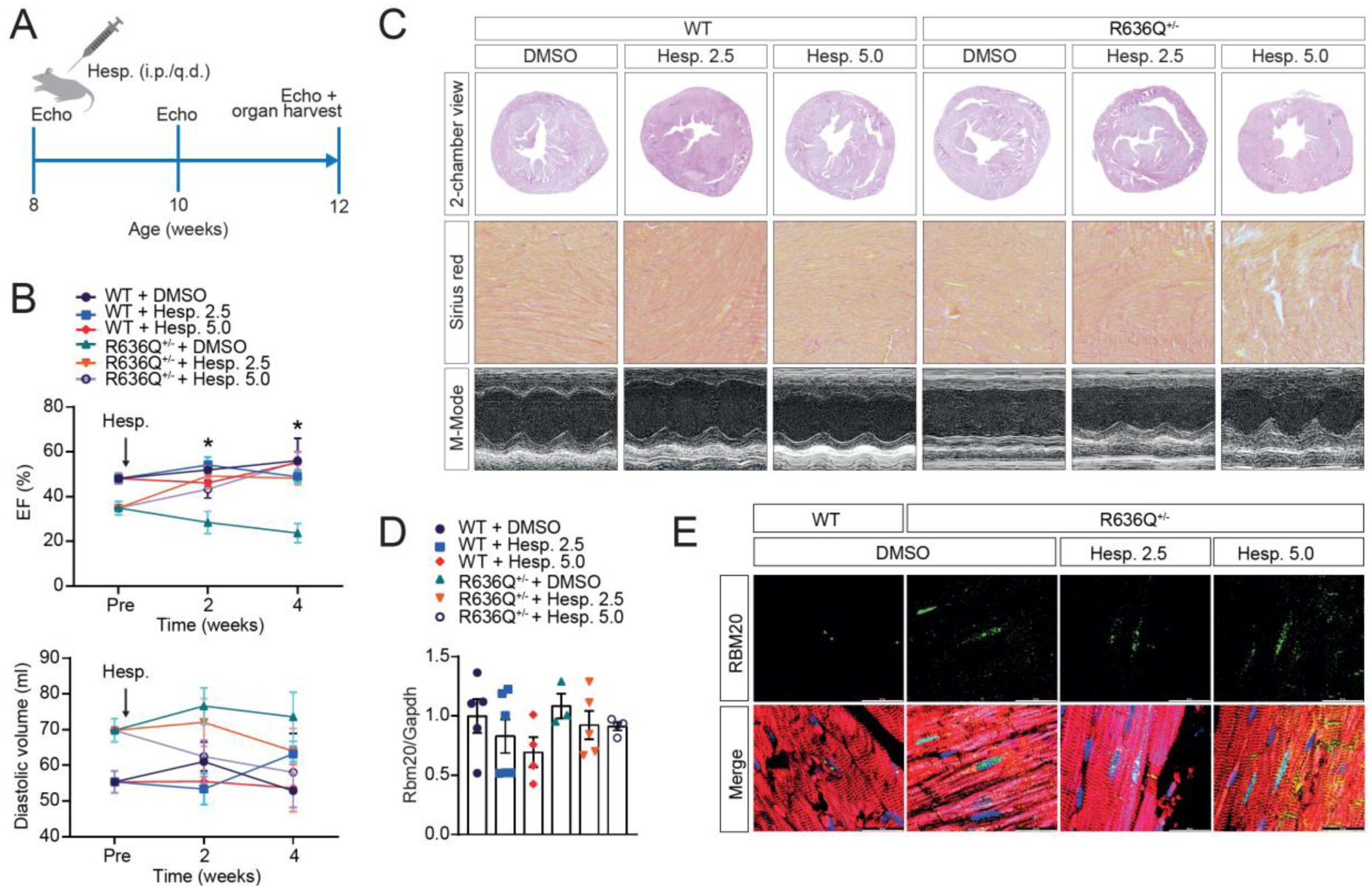
CAMK2D inhibition with hesperadin alleviates cardiac dysfunction in *Rbm20*-R636Q KI mice. A. Experimental set-up. B. Cardiac function and diastolic volume as measured by echocardiography. Significance was tested per time-point with a one-way ANOVA. C. H&E and Picrosirius Red staining. D. qPCR of *Rbm20*. H. Immunohistochemistry of RBM20 in left ventricular tissue.

### Hesperadin treatment inhibits CAMK2D activity in *Rbm20*-R636Q KI mice

We then performed RNA-sequencing on the hearts of these mice to investigate whether hesperadin treatment affected gene expression. We found similar numbers of differentially expressed genes (DEGs) in heterozygous *Rbm20*-R636Q KI mice treated with vehicle and with the two doses of hesperadin compared to WT hearts (Figure 5A). DEGs in the heterozygous *Rbm20*-R636Q KI mouse hearts included increased expression of cardiac stress markers such as *Anf*, and decreased expression of *Myh7b* (Figure 5B). The latter is in line with a previous observation of Ihara et al., who similarly found a loss of *Myh7b* in a Rbm20 KI mouse model [25]. We then overlapped DEGs in DMSO treated mice with DEGs in hesperadin treated mice, and found that only ∼55% of these DEGs were shared between groups, indicating that the transcriptional response differs between the DMSO and hesperadin treated mice (Figure 5C). Gene ontology enrichment of DEGs uniquely upregulated in heterozygous *Rbm20*-R636Q KI mouse hearts revealed an increase in calmodulin-dependent protein kinase activity in the heterozygous *Rbm20*-R636Q KI, and this enrichment was absent in the hesperadin treated heterozygous *Rbm20*-R636Q KI mouse hearts (Figure 5D, Supp Figure 7). Next, we performed phosphoproteomics, and found 782 phosphorylation events in 414 proteins that were increased in the hearts of heterozygous *Rbm20*-R636Q KI mice (Figure 5E). Kinase enrichment analysis showed that CAMK2D was one of the most enriched kinases predicted to be responsible for these phosphorylation events (Figure 5F). Among those, we then identified the phosphorylation events that were decreased with treatment of hesperadin. We found 365 events in 251 proteins, demonstrating that the phosphorylation of ∼half of these events was reversed (Cluster 1, Figure 5E). Moreover, kinase enrichment analysis of cluster 1 identified CAMK2D as one of the enriched kinases predicted to be responsible for these phosphorylation events (Figure 5G). Notably, AURKB, which is also inhibited by hesperadin, was likewise identified to be enriched in this cluster. Overall, this data shows that CAMK2D activity was increased in *Rbm20*-R636Q KI mice, and that hesperadin effectively inhibited this activity. Furthermore, our results suggest that pharmacological inhibition of CAMK2D is a promising therapeutic approach for patients with *RBM20* mutations.

**Figure 5.**
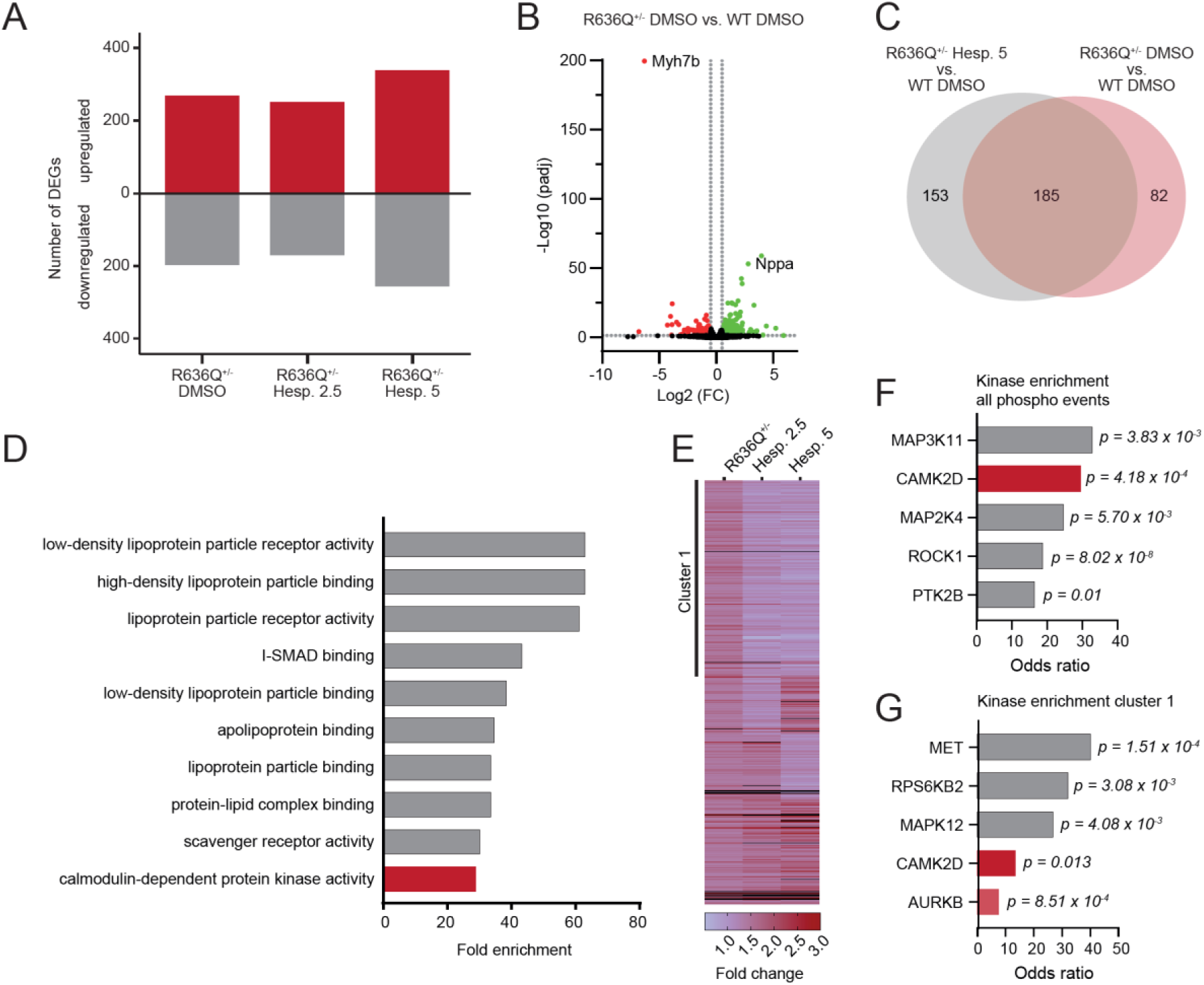
Hesperadin treatment inhibits CAMK2D activity in *Rbm20*-R636Q KI mice. A. Number of differentially expressed genes (DEGs) in vehicle and hesperadin treated heterozygous *Rbm20*-R636Q KI mice as compared to vehicle treated WT mice. B. Volcano plot of DEGs in heterozygous *Rbm20*- R636Q KI mouse hearts. C. Venn Diagram of upregulated genes in vehicle and hesperadin (5 mg/kg) treated heterozygous *Rbm20*-R636Q KI mouse hearts. D. Gene ontology enrichment analysis of genes uniquely upregulated in vehicle treated heterozygous *Rbm20*-R636Q KI mouse hearts. E. Heatmap of phosphorylation events in vehicle and hesperadin treated heterozygous *Rbm20*-R636Q KI mouse hearts. With fold change set to ≥1.3 we observed 782 events in 414 proteins. F. Kinase enrichment analysis of phosphorylation events increased (FC≥1.3) in heterozygous *Rbm20*-R636Q KI mouse hearts. G. Kinase enrichment analysis of phosphorylation events in Cluster 1 (365 events in 251 proteins).

## DISCUSSION

In this study, we investigated the relative contribution of CAMK2D to the detrimental phenotypes in RBM20 cardiomyopathy, and we established that heart failure in RBM20 cardiomyopathy depends on CAMK2D. Importantly, since TTN is misspliced in our models, this shows that RBM20 cardiomyopathy is not a TTN missplicing disease. AAV9-mediated re-expression of single *Camk2d* splice variants in *Rbm20*/*Camk2d* DKO mice re-introduces cardiac dysfunction irrespective of which splice isoform is re-expressed, unmasking that specific CAMK2D overactivation rather than missplicing underlies the severe phenotype. This also suggests there is considerable functional overlap between the different CAMK2D splice isoforms, at least in the setting of RBM20 cardiomyopathy. Multiple observations support this hypothesis: 1) the shift in CAMK2D splice isoform in mouse and human after loss of functional RBM20 is different, even though the cardiac phenotype is similar, 2) re-expression of all cardiac splice isoforms is sufficient to re-introduce cardiac dysfunction in *Rbm20/Camk2d* DKO mice, and 3) almost all CAMK2D protein was found in the nuclear fraction in both WT and *Rbm20* KO mouse hearts. Nevertheless, it will be important to further investigate the potential functional differences between the different splice isoforms of CAMK2D since AAV-mediated overexpression cannot precisely be controlled and variations in the amount and mosaic patterns may limit the interpretation of the approach that was used in this study. Importantly, we show that pharmacological inhibition of CAMK2D in *Rbm20*-R636Q KI mice has striking therapeutic effects on structural remodelling and cardiac dysfunction. Treatment of patients with *RBM20* mutations currently relies on the “fantastic four” (Betablockers, ACE inhibitors/AT1 antagonists, SGLT2 inhibitors and mineralocorticoid antagonists) [32]. As a specific approach, gene editing has been used in mouse models to correct disease causing mutations in *RBM20* [26, 33]. However, technical and safety hurdles with respect to successful AAV-mediated gene transfer and potentially CRISPR/CAS9 toxicity need to be overcome to translate this concept into the clinic. Moreover, this approach needs to be personalized due to the vast amount of different mutations that have been identified in the *RBM20* gene, which would require a costly individualized approach, limiting the feasibility. Here, we propose that pharmacological CAMK2D inhibition is a novel tailored approach to treat patients suffering from heart failure in *RBM20* cardiomyopathy with different mutations. It comes with great promise that after 35 years work on CAMK2 and several attempts to develop a specific CAMK2 inhibitor, a first inhibitor is in a phase II clinical trial (clinicaltrials.gov, identifier NCT06005428) [34, 35] . The use of hesperadin to inhibit CAMK2D in patients is not favourable because it also inhibits other kinases such as Aurora kinase B [31]. The current study, however, serves as a first proof of principle study that now calls for validation with a clinically tested CAMK2 inhibitor. Here, we used both a *Rbm20* KO model, which has only loss of splicing, and a *Rbm20* KI model, which has the additional detrimental cytoplasmic granule formation. In both cases, CAMK2D inhibition, either genetic in the KO model or pharmacological in the KI model, improved cardiac function. This suggests that pharmacological CAMK2D inhibition in RBM20 cardiomyopathy holds promise independent of the type of mutation in the patient, whether this is a purely loss-of-function variant (e.g. truncating variant) or a mislocalizing variant (e.g. RSRSP variant). Thus, CAMK2D inhibition can soon be tested as a first cause-directed DCM therapy. To do so, large cardiomyopathy registries have been established where the included patients undergo whole genome sequencing to identify the underlying disease mutations that could be selected as patients for a first tailored phase IIb/III trial [36]. Due to the here described specific CAMK2-dependent etiology of RBM20 cardiomyopathy, it is expected that a clinical trial would be sufficiently powered by the inclusion of a lower number of patients than in past clinical trials that included heart failure patients with heterogenous etiologies [37–40]. The development and success of mavacamten for HCM has demonstrated that only 251 patients, of which 123 were treated with mavacamten, were needed to demonstrate efficacy [7, 41]. Following this strategy pharmacological CAMK2 inhibition might become the first tailored DCM therapy on top on the general standard heart failure medication.

## Acknowledgements

The authors gratefully acknowledge Dr. Marco Hagenmüller for graphical assistance, Dr. Esther Creemers for supplying the *Rbm20* KO mouse line, Michaela Oestringer for coordination of the mouse lines, and Dr. Jennifer Schwarz and Dr. Frank Stein from the Proteomics Core at the European Molecular Biology Laboratory (EMBL) Heidelberg for help with phosphoproteomics. The authors gratefully acknowledge the data storage service SDS@hd supported by the Ministry of Science, Research and the Arts Baden-Württemberg (MWK) and the German Research Foundation (DFG) through grant INST 35/1503-1 FUGG.

## Author contributions

Conceptualization: MvdH, JB. Methodology: MvdH, JD, TBB, EK, LK, DL, JH, LSc, JK, TB, MD. Contributed materials: MK, LSt, CD. Funding acquisition: MvdH, JB. Supervision: MvdH, JB. Writing – original draft: MvdH, MD, JB. Writing – review and editing: all authors.

## MATERIALS AND METHODS

### Animal Experiments

This study adheres to the EU Directive (2010/63/EU) and received approval from the Institutional Animal Care and Use Committee at the Regierungspräsidium Karlsruhe, Germany (approval numbers G233/17 and G225/20). Hesperadin treatment in *Rbm20*-R636Q KI mice conformed to the EMBL guidelines for the Use of Animals in Experiments and was reviewed and approved by the Institutional Animal Care and Use Committee (IACUC). At the conclusion of the experiments, mice were euthanized via cervical dislocation. Global *Rbm20* knockout mice (FVB genetic background), global *Camk2d* knockout mice, and *Rbm20*-R636Q mice were previously generated [14–16].

*Rbm20*/*Camk2d* double knockout (DKO) were obtained by crossing *Rbm20* and *Camk2d* knockout mice. Mice were given normal chow and water ad libitum, and kept on a 12-hour light/dark cycle. Hesperadin (SelleckChem, Catalog No. S1529) was dissolved in 5% DMSO, 40% PEG300, 5% Tween 80, and 50% ddH2O. Mice were intraperitoneally injected daily with either vehicle (5% DMSO, 40% PEG300, 5% Tween 80, and 50% ddH2O), 2.5 mg/kg Hesperadin, or 5 mg/kg Hesperadin.

### Echocardiography and catecholamine challenge

Mice were anesthetized using 1–1.5% isoflurane (Baxter Deutschland GmbH, Germany). The depth of anesthesia was confirmed by assessing reflexes before conducting echocardiography, and the heart rate and respiratory rate were monitored continuously. To maintain body temperature, the mice were placed on a heating pad. Imaging was performed using the Vevo 2100 or 3100 Imaging System with an MS400 Transducer (FUJIFILM VisualSonics, Inc., Canada). B-mode and M-mode images of the short and long axis at the midpapillary muscle level were captured. Measurements of left-ventricular end-diastolic and end-systolic diameters were taken, and ejection fraction (EF) and fractional shortening (FS) were calculated using the VisualSonics VevoLab software’s LV trace tool with the Teicholz equation, based on at least five contraction cycles. Arrhythmia induction was done by intraperitoneal injection of 2 mg/kg epinephrine and 120 mg/kg caffeine after which mice were monitored for survival.

### RNA isolation and qPCR + RT-PCR

Total RNA was extracted from left ventricular mouse heart tissue using TRIzol™ Reagent (ThermoFisher, USA). cDNA was synthesized from 1 µg of RNA using the primaREVERSE RT-Kit (Steinbrenner Laborsysteme GmbH, Germany, SL-9540-1000), following the manufacturer’s instructions. RT-PCR was conducted with HOT FIREPol® DNA Polymerase (Solis Biodyne, Estonia) according to the manufacturer’s protocol, followed by gel electrophoresis on 1.5% agarose gels. RT-qPCR was carried out using PowerUp™ SYBR™ Green Master Mix (ThermoFisher, USA) on a Lightcycler 480 II system (Roche, USA), with each sample run in technical duplicates. Data was analyzed using LinRegPCR version 2018.030, and primer sequences are available in Supplementary Table S1.

### RNA-sequencing

Total RNA from left ventricular tissue of *Rbm20/Camk2d* KO mice was depleted from ribosomal RNA, polyA-enriched, fragmented, and paired-end sequenced at BGI (Hongkong, CN). Total RNA from left ventricular tissue from hesperadin treated mice was extracted and sequence libraries were prepared using SMART-Seq Total RNA pico input (Takara Bio) and sequenced in paired-end mode on AVITI (PE75, Element Biosciences). RNA-seq reads were aligned to mouse genome reference (mm10 GRCm38) using the Rsubread R package. Quality control of the reads was performed using Fastqcr and Rqc packages, while adapter trimming was conducted with Trimmomatic using standard parameters. For data processing and analysis, we employed a R-based pipeline utilizing BiocManager for package management, ShortRead for sequence manipulation and reading. Differential gene expression analysis was performed using DESeq2 [42]. Functional enrichment analysis of differentially expressed genes was conducted using the ClusterProfiler package to identify overrepresented biological pathways, molecular functions, and cellular components in the Gene Ontology (GO) database and relevant pathways in the KEGG database [43]. Splice junction counts were extracted from read aligments with Regtools with default parameters and annotated with respective transcript identifier, for expressed transcripts (mean TPM > 1) [44].

### Protein isolation and Western blotting

Tissue was homogenized in RIPA buffer (50mM Tris-HCl pH8, 150mM NaCl, 1% NP-40, 0.5% sodium deoxycholate, 0.1% SDS) supplemented with protease inhibitor cocktail (Roche) using a TissueLyser II (Qiagen). Protein concentration was measured using a BCA assay (Pierce). Western blotting was performed following standard protocols. Antibodies used were: anti-CAMK2 (611293, BD Biosciences), anti-GAPDH (MAB374, Sigma), anti-Flag (F1804, Sigma).

### Phosphoproteomics

For phosphoproteomics on the *Rbm20/Camk2d* DKO; mouse heart left ventricular tissue of 10-week-old was lysed in RIPA buffer supplemented with a protease inhibitor (S8820, MilliporeSigma) and a phosphatase inhibitor (P5726, MilliporeSigma). Proteins were digested using Lysyl Endopeptidase (Lys-C, FUJIFILM Wako Pure Chemicals, Japan) and trypsin (MilliporeSigma). Desalting was carried out using Sep-Pak C18 columns (Waters), preconditioned with 100% acetonitrile (ACN) and rinsed twice with 0.1% trifluoroacetic acid (TFA). The samples were eluted in 60% ACN-0.1% TFA, then dried. Phosphopeptides were enriched using the High-Select TiO2 Phosphopeptide Enrichment Kit (A32993, Thermo Fisher) according to the manufacturer’s instructions, and the enriched samples were analyzed via mass spectrometry. Proteomic analysis employed an Easy nLC 1000 UHPLC system coupled to a Q Exactive Plus mass spectrometer (Thermo Fisher). Peptides were dissolved in Solvent A (0.1% FA), loaded with an autosampler, and separated on custom-made 50 cm fused silica columns [75-μm internal diameter (I.D.), packed with 2.7-μm C18 Poroshell beads (Agilent)] at a flow rate of 0.75 µL/min. Gradients of 90 minutes for phosphoproteomics and 240 minutes for whole proteome analysis were applied, with peptides eluted at 250 nL/min. The eluted peptides were ionized via nanoelectrospray and introduced into the mass spectrometer’s heated transfer capillary. The instrument was operated in a data-dependent acquisition mode. Full MS scans (300–1,750 m/z) were performed in the Orbitrap at a resolution of 70,000, with an automated gain control (AGC) target of 3 × 10⁶ ions and a maximum injection time of 20 ms. A dynamic exclusion window of 20 seconds was used, and the top 10 most intense peaks (z ≥ 2) from each scan were fragmented in the high-energy collision-induced dissociation (HCD) cell with a normalized collision energy of 25%. MS/MS scans were acquired at a resolution of 17,500. Data were processed using MaxQuant software and its integrated Andromeda search engine, with default settings and Phospho (STY) included as a variable modification. Statistical analyses were conducted using Perseus.

For phosphoproteomics on hesperadin treated *Rbm20*-R636Q^+/-^ mice; sample preparation for phosphoproteome analysis was performed essentially as described by Potel et al. [45]. Briefly, mouse heart left ventricular tissue was lysed in 500 µL of lysis buffer consisting of 100 mM Tris-HCl (pH 8.5), 7 M urea, 1% Triton X-100, 5 mM Tris(2-carboxyethyl)phosphine hydrochloride (TCEP), 55 mM 2-chloroacetamide (CAA), 10 U/mL DNase I, 1 mM magnesium chloride, 1 mM sodium orthovanadate, Phosphatase Inhibitor Cocktail 2 (Sigma Aldrich), and Halt Protease Inhibitor Cocktail (Thermo Fisher). The lysates were sonicated 45 minutes (20 seconds on, 40 seconds off) at 4°C, using a Bioruptor (Diagenode). Cell debris was removed by centrifugation at 14,000 × g for 10 minutes at 4°C. Benzonase (final concentration 1%) was added to the cleared supernatant, and the mixture was incubated at room temperature for 30 minutes. Protein concentrations were determined using a Bradford assay. For each sample, 500 µg of protein was precipitated by adding sequentially four volumes of methanol, one volume of chloroform, and three volumes of ultrapure water. After centrifugation at 4,000 × g for 15 minutes, the upper aqueous phase was removed, and three volumes of methanol were added, followed by an additional centrifugation step. The liquid phase was discarded, and the resulting protein pellet was air-dried. The pellet was subsequently resuspended in 50 mM HEPES/NaOH (pH 8.5), 1% sodium deoxycholate (SDC), 5 mM TCEP, and 30 mM CAA. Proteins were digested overnight at room temperature with trypsin added at a protein-to-enzyme ratio of 50:1. Digestion was stopped by the addition of trifluoroacetic acid (TFA) to a final concentration of 1% in the sample. The sodium deoxycholate was precipitated for 15 minutes at room temperature and the samples were centrifuged at 17,000 × g for 10 minutes at room temperature. Peptides were desalted using Oasis HLB 96-well plates (30 µm, Waters) under gravity flow. Washing was performed with buffer A (MS-grade water, 0.1% formic acid) and peptides were eluted using buffer B (80% acetonitrile, 0.1% formic acid). Eluted peptides were dried by vacuum centrifugation and stored at -80°C until phosphopeptide enrichment. Phosphopeptide enrichment was performed essentially as previously described by Leutert et al. [46]. Briefly, peptides were resuspended in IMAC loading solvent (80% acetonitrile, 0.4% TFA). Of each sample a small aliquot was used for full proteome analysis, while the remaining peptides were subjected to phosphopeptide enrichment using the KingFisher Apex™ platform (Thermo Fisher) and magnetic Fe- NTA beads (Cube Biotech). Enriched phosphopeptides were eluted with 0.2% dimethylamine in 80% acetonitrile to facilitate subsequent TMT labeling. Peptides were modified with TMTpro labelling reagent (Thermo Fisher Scientific) [47]. In short, 0.5mg reagent was dissolved in 45ul acetonitrile (100%) and 8 ul of stock was added and incubated for 1h room temperature. Followed by quenching the reaction with 5% hydroxylamine for 15min. RT. Samples were combined for multiplexing and an OASIS® HLB µElution Plate (Waters) was used for further sample clean up. Offline high pH reverse phase fractionation was performed using an Agilent 1200 Infinity high-performance liquid chromatography (HPLC) system equipped with a quaternary pump, degasser, variable wavelength UV detector (set to 254 nm), peltier-cooled autosampler, and fraction collector (both set at 10°C for all samples). The column was a Gemini C18 column (3 μm, 110 Å, 100 x 1.0 mm, Phenomenex) with a Gemini C18, 4 x 2.0 mm SecurityGuard (Phenomenex) cartridge as a guard column. The solvent system consisted of 20 mM ammonium formate (pH 10.0) (A) and 100% acetonitrile as mobile phase (B). The separation was accomplished at a mobile phase flow rate of 0.1 mL/min using the following linear gradient: 100% A for 2 min., from 100% A to 35% B in 59 min., to 85% B in a further 1 min., and held at 85% B for an additional 15 min., before returning to 100% A and re-equilibration for 13 min. Forty-eight fractions were collected along with the LC separation that were subsequently pooled into 12. Whereby the first fraction was not used at all. Pooled fractions were dried under vacuum centrifugation. An UltiMate 3000 RSLC nano LC system (Dionex) fitted with a trapping cartridge (µ-Precolumn C18 PepMap 100, 5µm, 300 µm i.d. x 5 mm, 100 Å, Thermo Fisher Scientific) and an analytical column (nanoEase™ M/Z HSS T3 column 75 µm x 250 mm C18, 1.8 µm, 100 Å, Waters). Trapping was carried out with a constant flow of trapping solution (0.05% trifluoroacetic acid in water) at 30 µL/min onto the trapping column for 6 minutes. Subsequently, peptides were eluted via the analytical column running solvent A (3%DMSO, 0.1% formic acid in water) with a constant flow of 0.3 µL/min, with increasing percentage of solvent B (3% DMSO, 0.1% formic acid in acetonitrile). Each fraction of the full proteome was measured for 120 min and each fraction of the phopshoproteme for 90 min. The peptides were introduced into a Orbitrap Fusion™ Lumos™ Tribrid™ Mass Spectrometer via a Pico-Tip Emitter 360 µm OD x 20 µm ID; 10 µm tip (CoAnn Technologies) and an applied spray voltage of 2.2 kV. The capillary temperature was set at 275°C. Full mass scan was acquired with mass range 375-1500 m/z (to 1650 for phosphopeptides) in profile mode in the orbitrap with resolution of 120000. The filling time was set at maximum of 50 ms with an AGC target set to standard which allows absolute AGC target of 4x105 ions. Data dependent acquisition (DDA) was performed with the resolution of the Orbitrap set to 30000, with a fill time of 94 ms (110 ms for phoshopeptides) and a limitation of 1x105 ions. A normalized collision energy of 34 was applied. MS2 data was acquired in profile mode. The isolation window of the quadrupole was set to 0.7 m/z. Dynamic exclusion time of 60 s (20s for the phosphopeptides) was used. Fragpipe v22.1 with MSFragger v4.1 was used to process the acquired data, which was searched against the mus musculus Uniprot proteome database (UP000000589, ID10090, 21986 entries, release October 2022) with common contaminants and reversed sequences included [48]. The following modifications were considered as fixed modification: Carbamidomethyl (C) and TMT6 or 10 (K). As variable modifiations: Acetyl (Protein N-term), Oxidation (M) and TMT6 or 10 (N-term), for the phosphoproteome specifically phosphorylation on STY. For the MS1 and MS2 scans a mass error tolerance of 20 ppm was set. Further parameters were: Trypsin as protease with an allowance of maximum two missed cleavages; Minimum peptide length of seven amino acids; at least two unique peptides were required for a protein identification. The false discovery rate on peptide and protein level was set to 0.01. Data was then analyzed as follows: the raw output files of FragPipe (psm.tsv for phospho data and protein.tsv files for input data) were processed using the R programming language (ISBN 3-900051-07-0). Only peptide spectral matches (PSMs) with a phosphorylation probability greater 0.5 and proteins with at least 2 unique peptides were considered for the analysis.

Phosphorylated amino acids were marked with a * in the amino acid sequences behind the phosphorylated amino acid, labeled with a 1, 2 or 3 for the number of phosphorylation sites in the peptide and concatenated with the protein ID in order to create a unique ID for each phosphopeptide. Raw TMT reporter ion intensities were summed for all PSMs with the same phosphopeptide ID. For the input data, the reporter ion intensities were used as given in the protein.tsv output files. Phospho signals were also normalized by input abundance. For this the reporter ion intensity for each unique phosphor ID, condition and replicate was normalized according to the following formula:

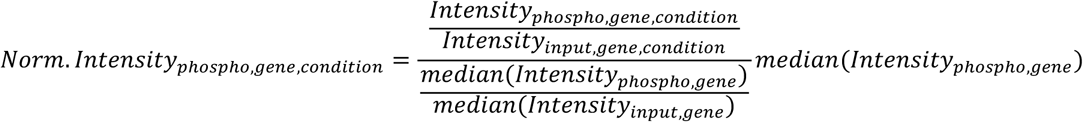

Transformed summed TMT reporter ion intensities were first cleaned for batch effects using the ‘removeBatchEffects’ function of the limma package and further normalized using the vsn package (variance stabilization normalization)[49, 50]. Missing values were imputed with ‘knn’ method using the Msnbase package [51]. Proteins were tested for differential expression using the limma package. Phospho, Normalized phospho and input data were tested separately. The replicate information was added as a factor in the design matrix given as an argument to the ‘lmFit’ function of limma. Also, imputed values were given a weight of 0.05 in the ‘lmFit’ function. A phosphopeptide or protein was annotated as a hit with a false discovery rate (fdr) smaller 5 % and a fold-change of at least 100 % and as a candidate with a fdr below 20 % and a fold-change of at least 50 %. Clustering with all hit peptides based on the median abundances normalized by median of control condition was conducted to identify groups of peptides with similar patterns across conditions. The ’kmeans’ method was employed, using Euclidean distance as the distance metric and ’ward.D2’ linkage for hierarchical clustering. The optimal number of clusters (10) was determined using the Elbow method, which identifies the point where the within-group sum of squares stabilizes. Gene ontology (GO) enrichment analysis was performed using the ’compareCluster’ function of the ’clusterProfiler’ package, which assesses over-representation of GO terms in the dataset relative to the background gene set [52]. Enrichment was conducted for the following GO categories: Cellular Component (CC), Molecular Function (MF), and Biological Process (BP). The analysis was performed using ’org.Mm.eg.db’ as the reference database. The odds ratio (’odds_ratio’) for each GO term was calculated by comparing the proportion of genes associated with that term in the dataset (’GeneRatio’) to the proportion in the background set (’BgRatio’). An odds ratio greater than 1 indicates that the GO term is enriched in the dataset compared to the expected background. Kinase enrichment analysis was done using KEA3 [53].

### Histochemistry

After overnight fixation in 4% PFA, mouse hearts were embedded in paraffin using the HistoCore PEARL Tissue Processor and Arcadia Embedding Center (both from Leica Biosystems, Germany). Sections 5 µm thick were cut, mounted on slides, and dried on a hotplate for 30 minutes, followed by overnight drying in a 37°C incubator. The sections were dewaxed twice in ROTI®Histol (Carl Roth GmbH + Co. KG, Germany) for 10 minutes each, then rehydrated through a descending alcohol series (100% for 2 minutes, 96% for 2 minutes, and 70% for 2 minutes) before being placed in distilled water. For HE staining, an H&E fast staining kit (Carl Roth GmbH + Co. KG, Germany) was used according to the manufacturer’s instructions. Sirius Red staining was performed by immersing slides in Picro-Sirius Red Solution (ScyTek Laboratories, USA) for 30 minutes, followed by differentiation in acidified water (1% acetic acid) twice. The slides were then dehydrated using an ascending alcohol series (70% for 2 minutes, 96% for 2 minutes, and 100% for 2 minutes), cleared in ROTI®Histol, and mounted with 35 µl EUKITT® neo mounting medium (ORSAtec GmbH, Germany). The slides were scanned using a ZEISS Axioscan slide scanner (ZEISS, Germany).

### Immunohistochemistry

Left ventricular sections were deparaffinized, rehydrated in a series of ethanol, and boiled for 5 min in antigen unmasking solution (10mM Citrate, pH 6) in a pressure-cooker. Sections were subsequently incubated with 4% normal goat serum in PBS with 0.1% Triton X-100 for 60 min at RT, and incubated with primary antibody in 4% normal goat serum in PBS overnight at 4° C. Primary antibodies used were: anti-RBM20 (PA5-58068, Thermo Fisher) and anti-alpha actinin (A7811, Sigma). Alexa Fluor 488 and Alexa Fluor 594 conjugated antibodies (Invitrogen) were used as secondary antibodies, and nuclei were visualized using DAPI (Sigma). Pictures were taken on a confocal microscope (Leica MICA, Leica Microsystems, Germany).

### AAV generation and injection

The *Camk2d* splice variants were amplified from cDNA of mouse cardiac tissue, N-terminally FLAG- tagged and cloned into a p-GEM-T vector (Promega) and subsequently shuttled into the AAV genome-plasmid pSSV9-TnT (kindly provided by Prof. Oliver Müller [54]) containing the cardiac- specific human troponin-T promotor, a chimeric human ß-gobin/immunoglobulin heavy chain intron and a SV40 polyadenylation signal. These constructs and a control humanized Renilla luciferase encoding plasmid, pSSV9-TnT-hrLuc, together with the adeno-helper plasmid, pdp9rs (kindly provided by Dr. J. Kleinschmidt), were used to generate single-stranded, pseudotyped ssAAV2/9 in the AAVpro® 293T Cell Line (Takara) as described previously [55]. In brief, each AAV genome plasmid was co-transfected with the adeno-helper plasmid using Polyethylenimine (PEI). 72 hours after transfection the supernatants and cells were harvested. The first underwent ammonium-sulfate precipitation, while the later was lysed by freeze-thawing followed by DNase digestion. AAV vectors were then purified by ultracentrifugation on iodixanol step gradients, buffer exchanged to PBS, pooled and concentrated. The final vector stocks were quantified by qPCR using a plasmid standard curve. Mice were briefly anesthetized and injected retro-orbitally with 5 x 10^12^ viral genomes.

### Statistical analysis

Data are presented as mean ± sem, and were analyzed with appropriate statistical tests, as indicated in the respective figure legends. A value of p < 0.05 was considered statistically significant.

---

**Supplemental Figure 1.**
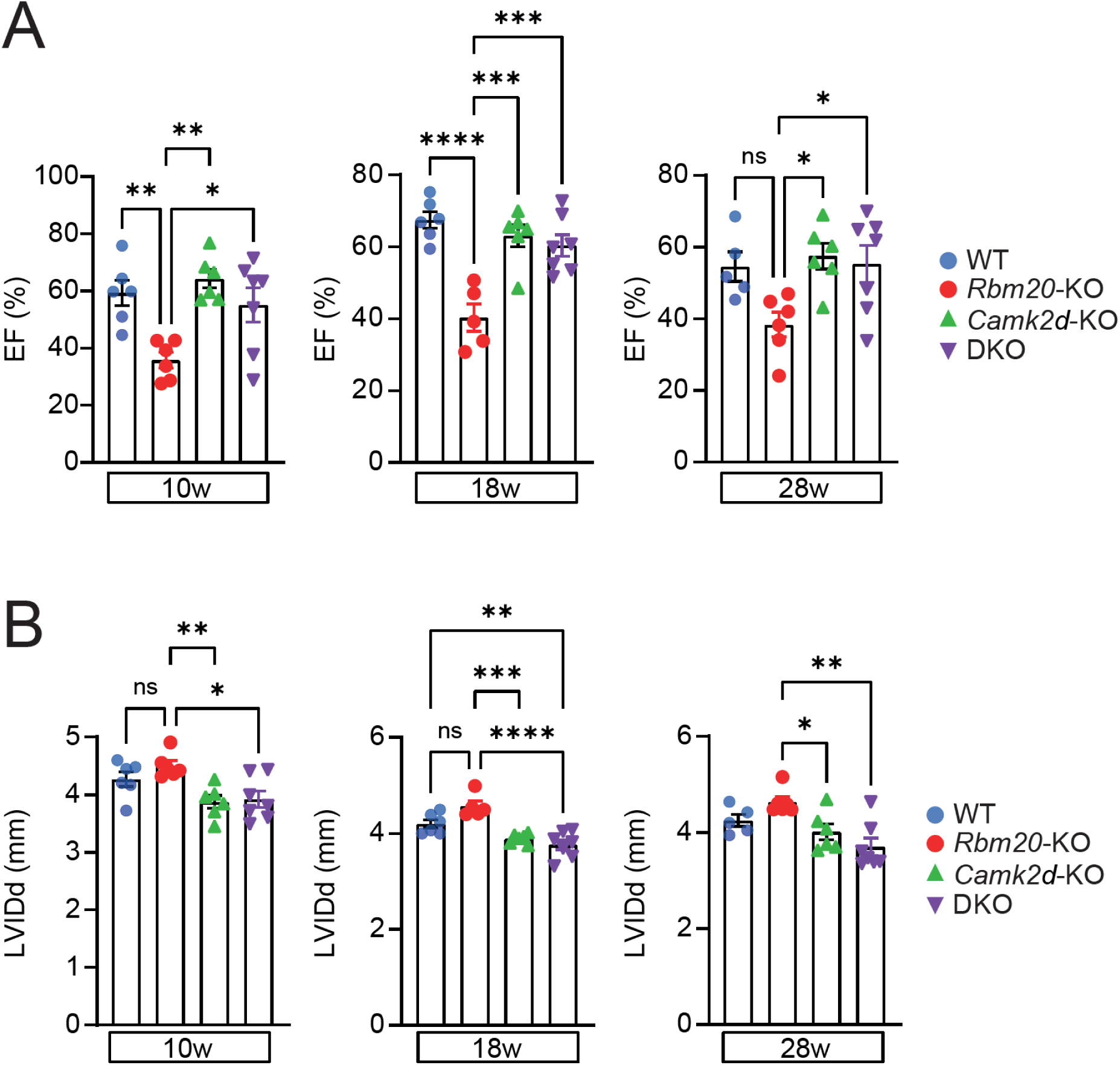
Echocardiography of *Rbm20/Camk2d* DKO mice. A. Ejection fraction (EF) as measured by echocardiography at 10, 18, and 28 weeks of age. B. Left ventricular internal diameter at diastole (LVIDd) as measured by echocardiography at 10, 18, and 28 weeks of age. Significance was tested with a one-way ANOVA.

**Supplemental Figure 2.**
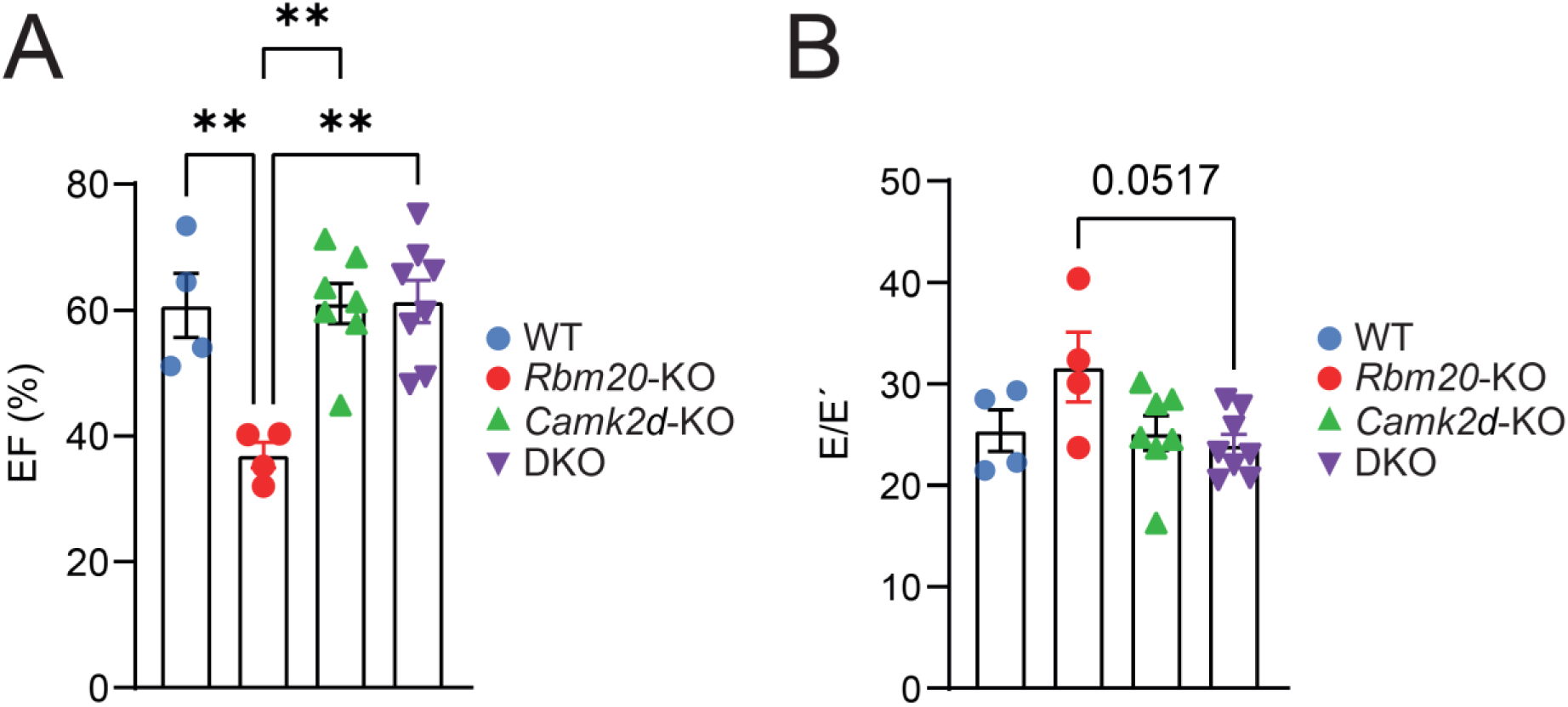
Diastolic function of *Rbm20/Camk2d* DKO mice. A. Ejection fraction (EF) as measured by echocardiography at 12 weeks of age. B. E/E’ ration as measured by echocardiography at 12 weeks of age. Significance was tested with a one-way ANOVA.

**Supplemental Figure 3.**
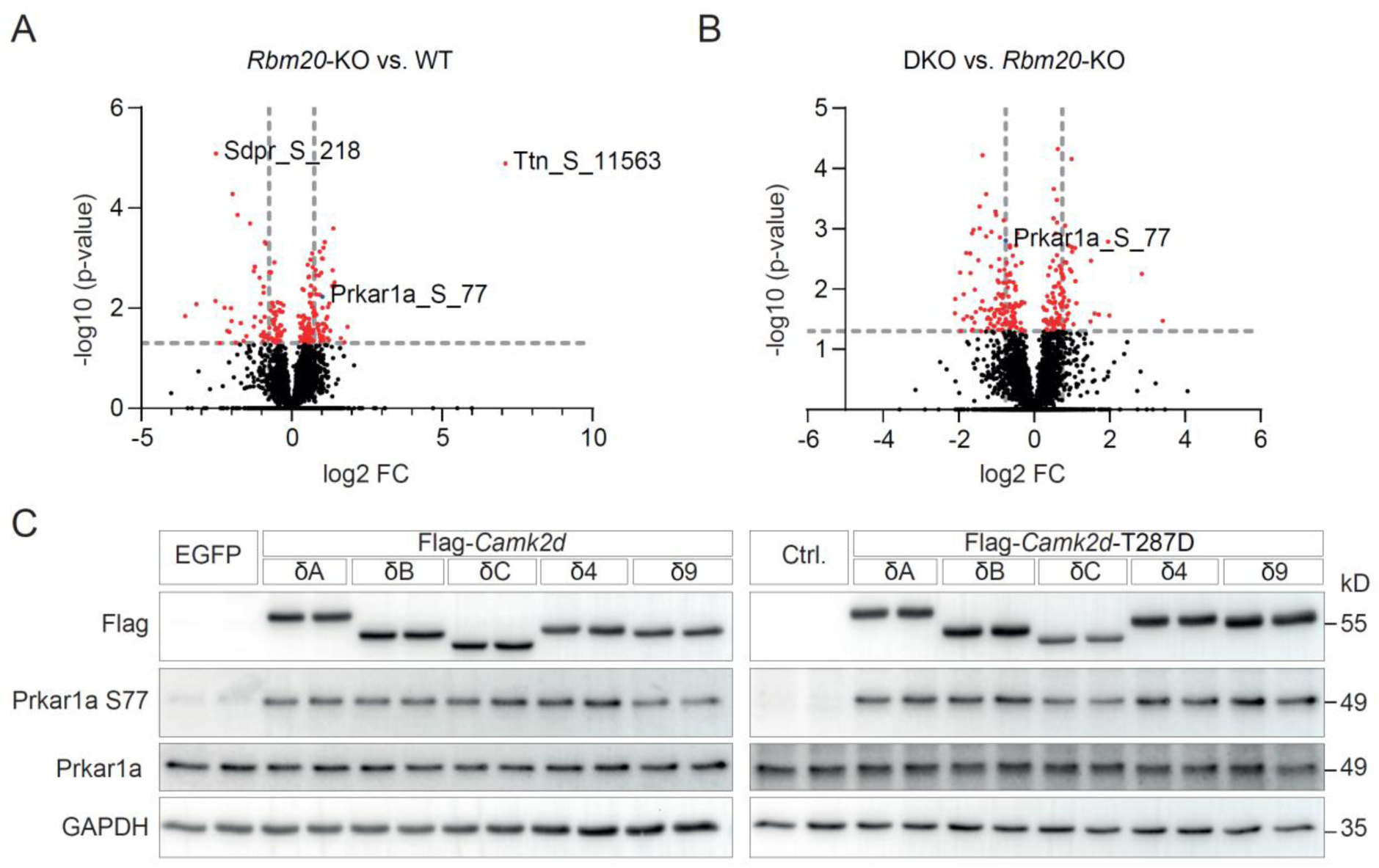
Phosphoproteomics of *Rbm20/Camk2d* DKO mouse hearts. A. Volcano plot of differentially phosphorylated proteins in the hearts of *Rbm20* KO mice. B. Volcano plot of differentially phosphorylated proteins in the hearts of DKO vs. *Rbm20* KO mice. C. Overexpression of WT and constitutively active (T287D) CAMK2D splice variants in HEK293 cells is sufficient to induce PRKAR1A phosphorylation at Serine 77.

**Supplemental Figure 4.**
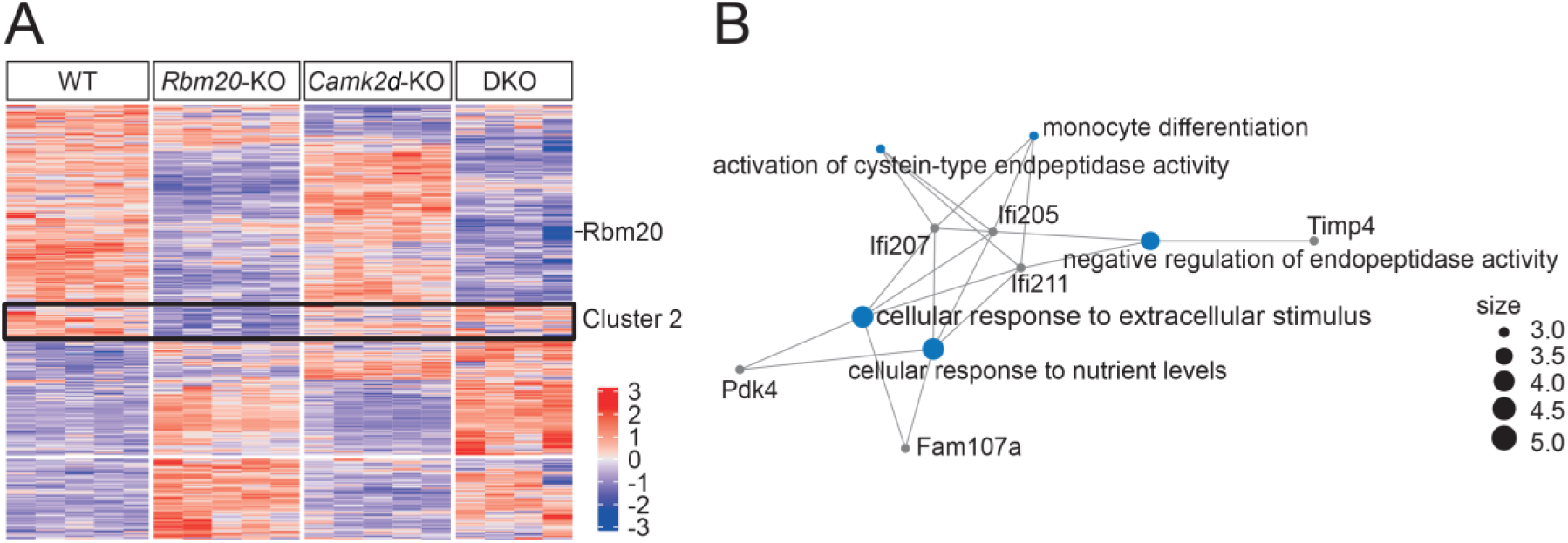
Transcriptomic analysis of *Rbm20/Camk2d* DKO mouse hearts. A. Heatmap of DEGs in the hearts of WT, *Rbm20* KO, *Camk2d* KO, and *Rbm20/Camk2d* DKO mice. B. Gene set enrichment analysis of cluster 2.

**Supplemental Figure 5.**
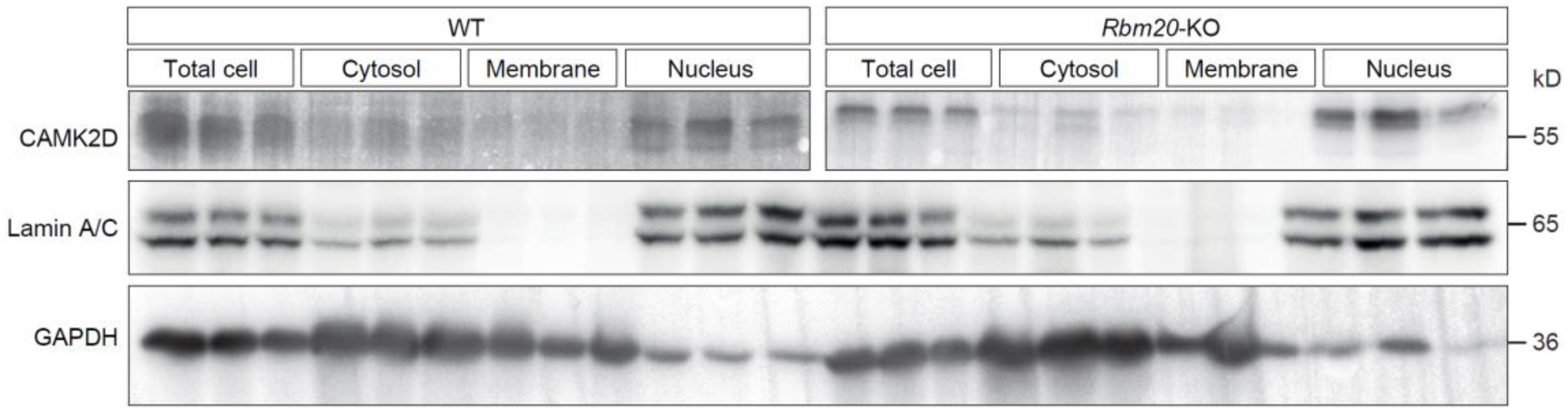
Fractionation of left ventricular tissue of WT and *Rbm20* KO mice. Cellular fractionation of left ventricular tissue of WT and *Rbm20* KO mouse hearts. Lamin A/C denotes the nuclear fraction, while GAPDH denotes the cytoplasmic fraction.

**Supplemental Figure 6.**
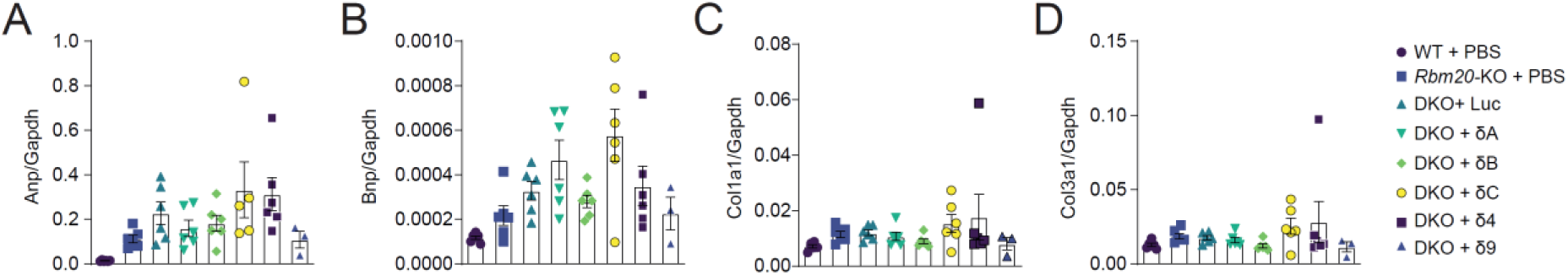
Expression of marker genes in DKO mouse hearts with re-expression of CAMK2D. A-D. qPCR analysis of *Anp* (*Anf*), *Bnp*, *Col1a1*, and *Col3a1* in the hearts of DKO mice after re-expression of single cardiac CAMK2D splice variants.

**Supplemental Figure 7.**
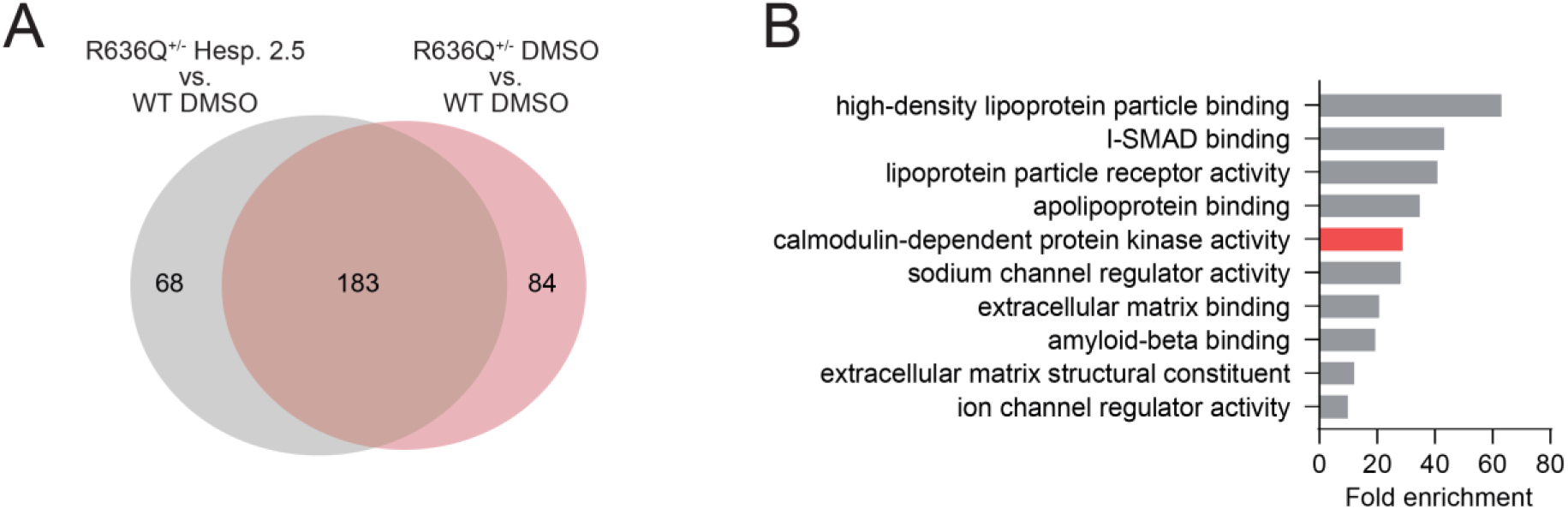
Hesperadin treatment inhibits CAMK2D activity in *Rbm20*-R636Q KI mice. A. Venn Diagram of upregulated genes in vehicle and hesperadin (2.5 mg/kg) treated heterozygous *Rbm20*-R636Q KI mouse hearts. B. Gene ontology enrichment analysis of genes uniquely upregulated in vehicle treated heterozygous *Rbm20*-R636Q KI mouse hearts.

**Supplementary Table S1:**
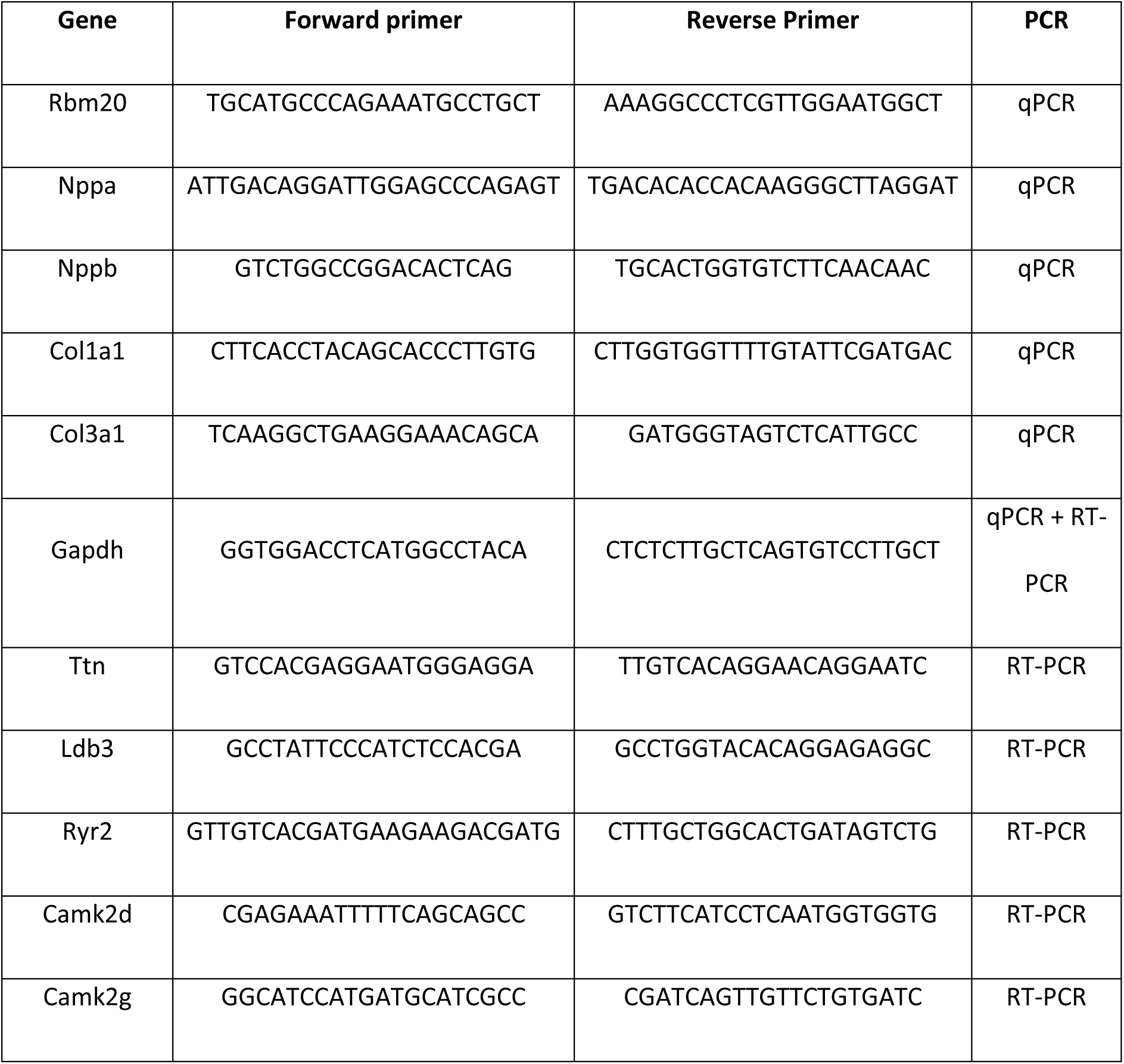
Primer sequences.

**Supplemental Table S2:** Software versions.

| Software: | Version: | DOI |
| --- | --- | --- |
| R | 4.2.3 |  |
| DESeq2 | 1.38.3 | 10.18129/B9.bioc.DESeq2 |
| tximport | 1.26.1 | 10.18129/B9.bioc.tximport |
| ComplexHeatmap | 2.12.1 | 10.18129/B9.bioc.ComplexHeatmap |
| clusterProfiler | 4.4.4 | [10.18129/B9.bioc.clusterProfiler](https://doi.org/10.18129/B9.bioc.clusterProfiler) |
| regtools | 1.0.0 | 10.1038/s41467-023-37266-6 |

